# MitoWave: Spatio-temporal analysis of mitochondrial membrane potential fluctuations during ischemia-reperfusion

**DOI:** 10.1101/2020.05.21.108670

**Authors:** D. Ashok, B. O’Rourke

## Abstract

Mitochondria exhibit non-stationary unstable membrane potential (ΔΨ_m_) when subjected to stress, such as during Ischemia/Reperfusion (I/R). Understanding the mechanism of ΔΨ_m_ instability involves characterizing and quantifying this phenomenon in response to I/R stress in an unbiased and reproducible manner. We designed a simple ImageJ-MATLAB-based workflow called ‘MitoWave’ to unravel dynamic mitochondrial ΔΨ_m_ changes that occur during ischemia and reperfusion. MitoWave employs MATLAB’s wavelet transform toolbox. *In-vitro* Ischemia was effected by placing a glass coverslip for 60 minutes on a monolayer of neonatal mouse ventricular myocytes (NMVMs). Removal of the coverslip allowed for reperfusion. ΔΨ_m_ response to I/R was recorded on a confocal microscope using TMRM as the indicator. As proof-of-principle, we used MitoWave analysis on ten invitro I/R experiments. Visual observations corroborated quantitative MitoWave analysis results in classifying the ten I/R experiments into five outcomes that were observed based on the oscillatory state of ΔΨ_m_ throughout the reperfusion time period. Statistical analysis of the distribution of oscillating mitochondrial clusters during reperfusion shows significant differences between five different outcomes (p< 0.001). Features such as time-points of ΔΨ_m_ depolarization during I/R, area of mitochondrial clusters and time-resolved frequency components during reperfusion were determined per cell and per mitochondrial cluster. We found that mitochondria from NMVMs subjected to I/R oscillate in the frequency range of 8.6-45mHz, with a mean of 8.73±4.35mHz. Oscillating clusters had smaller areas ranging from 49.78±40.64 μm^2^ while non-oscillating clusters had larger areas 65.97±42.07μm^2^. A negative correlation between frequency and mitochondrial cluster area was seen. We also observed that late ΔΨ_m_ loss during ischemia correlated with early ΔΨ_m_ stabilization after oscillation on reperfusion. Thus, MitoWave analysis provides a way to quantify complex time-resolved mitochondrial behavior. It provides an easy to follow workflow to automate microscopy analysis and allows for unbiased, reproducible quantitation of complex nonstationary cellular phenomena.

**Statement of Significance:** Understanding mitochondrial instability in Ischemia Reperfusion injury is key to determining efficacy of interventions. The MitoWave analysis is a powerful yet simple tool that enables even beginner MATALAB-Image J users to automate analysis of time-series from microscopy data. While we used it to detect ΔΨ_m_ changes during I/R, it can be adapted to detect any such spatio-temporal changes. It standardizes the quantitative analysis of complex biological signals, opens the door to in-depth screening of the genes, proteins and mechanisms underlying metabolic recovery after ischemia-reperfusion.

## Introduction

Spatio-temporal oscillations (electrical and contractile) are fundamental to normal cardiac function but are also a potential source of pathological instability and chaos (1). A stable supply of energy is required to prevent maladaptive emergent phenomena, and mitochondria are well-suited to dynamically adapt to the varying workloads of the organism. Nevertheless, both under physiological conditions (2) or after metabolic stress, mitochondrial oscillations (3), flickers(4),(5), transients(6), or fluctuations(7)(8) have been observed, when parameters such as ΔΨ_m_, flavin or NADH redox potential, pH, or Reactive Oxygen Species (ROS) have been measured. For example, ΔΨ_m_, ROS and NADH were shown to oscillate in a self-sustaining manner in adult cardiomyocytes subjected to substrate deprivation (9) or oxidative stress (10) in a frequency range spanning from ~1-40 mHz (11). Similarly, local mitochondrial superoxide oscillations (“mitoflashes”) in cardiomyocytes had a frequency of ~40mHz (12). As we have previously reported, ΔΨ_m_ oscillation also reproducibly occurs upon reperfusion after ischemia in neonatal rat ventricular myocyte monolayers (13). Importantly, interventions that suppressed mitochondrial ΔΨ_m_ instability on reperfusion also abrogated cardiac arrhythmias, both in neonatal myocytes (13) and isolated perfused hearts(14),(15). Hence, understanding the mechanism of mitochondrial destabilization during oxidative stress or ischemia/reperfusion (I/R) injury is essential to develop novel therapeutic strategies to prevent cardiac arrhythmias and contractile dysfunction associated with metabolic stress.

Determining the efficacy of interventions targeting spatiotemporal changes in mitochondria requires a robust, unbiased analytical approach, yet there are few reports describing methods for the automated analysis of non-stationary fluctuations observed in image time series. We have previously employed wavelet transform as a tool for characterizing ΔΨ_m_ oscillations and to describe dynamic mitochondrial clustering in adult cardiac myocytes by employing a mesh grid to outline individual mitochondrial clusters (16)(17). Here, we describe a workflow for characterizing spatially distributed ΔΨm loss and oscillation during I/R in terms of time-resolved frequency components, area of mitochondrial clusters, and times of reversible (ischemia) or irreversible (reperfusion) ΔΨm loss in neonatal cardiac cell monolayers. We apply discrete or continuous wavelet transform methods, followed by feature extraction, to analyze reperfusion-induced unsynchronized ΔΨm oscillations in neonatal ventricular myocytes. The method accurately identifies key transitions in mitochondrial behavior during I/R and quantifies the principal frequency components of mitochondrial instability and how they evolve over time. Moreover, the method is generalizable to the analysis of spatiotemporal variation of any parameter recorded during image time series. The method provides a workflow to automate microscopy analysis and allows for unbiased, reproducible quantitation of complex nonstationary cellular phenomena.

## Methods

### Neonatal cardiomyocyte isolation and cell culture

Neonatal mouse cardiac myocytes (NMCMs) were isolated using the MACS cell separation kit (Miltenyi Biotec: Catalog #130-100-825 and #130-098-373). Briefly, hearts from 0-2 day old mice were excised, chopped into small pieces and digested using reagents supplied by the kit. A cardiomyocyte-rich cell suspension was obtained by separation of magnetically labelled non-cardiac cells from total cell suspension upon application of a magnetic field. 1X10^6^ NMCMs were plated on fibronectin-coated (10μg/ml) 35mm (D=20mm) glass coverslip dishes (NEST^®^ catalog # 801001) in Medium-199 supplemented with 25mM HEPES, 2μg/ml Vitamin B12, 50U/ml Pen-strep, 1X non-essential 286 Amino acids and 10% FBS. The next day, the medium was changed to 2% FBS medium. Ischemia/Reperfusion experiments and imaging were performed on the 5th-6th day of culture.

### Inducing Ischemia and Reperfusion and ΔΨ_m_ Imaging

To monitor mitochondrial inner membrane potential (ΔΨ_m_), 50nM Tetramethylrhodamine methylester (TMRM) was loaded for 30 min at 37°C prior to the start of the experiment and the media was then replaced with fresh Tyrode's buffer (130mM NaCl, 5mM KCl, 1mM MgCl_2_, 10mM NaHEPES, 1mM CaCl_2_ and 5mM Glucose). Experiment was performed at 37°C. A typical protocol included a baseline reading for 10 minutes followed by 60 minutes of regional ischemia induced by placing a glass coverslip and followed by 60 minutes of reperfusion upon removal of the coverslip, as previously described in neonatal rat ventricular myocytes (13),(18). During this 130-minute period, images were obtained every 15 sec on a laser-scanning confocal microscope (Olympus FV3000RS). TMRM fluorescence was imaged using a 40X silicone-immersion objective (Olympus UPLSAPO40XS) with 561nm excitation/ 570-620nm emission. Cells were imaged in Galvano scanning mode without averaging. Each image was 16-bit with a size of 318.2X319.2 microns (512X512 pixels). To minimize laser-induced damage during the long protocol, a neutral density filter of 10% was applied in the excitation path and the laser intensity was set by the software to 0.06% power (20 mW 561 nm LED laser). At the 15 sec image acquisition interval, only frequencies below 66.67mHz are resolvable based on Nyquist–Shannon sampling theorem (19).

### Image Analysis

Image series of the time-course of Ischemia/Reperfusion experiments were analyzed using Fiji (https://imagej.net/Fiji/Downloads). A custom-built segmentation-analysis macro was generated to track each cell’s ΔΨ_m_ during the in-vitro I/R injury. ΔΨ_m_ response to I/R was analyzed at the cellular level by segmentation analysis (ImageJ). Steps for segmentation analysis included a pre-processing step to align the images in the stack using a ‘StackReg’ plugin (20). Segmentation of each cell was done by applying a median filter (radius=2) to the first image of the stack and then applying an auto local threshold (Niblack). All particles above the radius of 60 were included in the analysis. TMRM fluorescence intensity for each cell over Ischemia and Reperfusion were obtained. See supplement for macros.

### Discrete and Continuous wavelet transform

Limited information can be obtained through the use of frequency domain methods such as Fourier transform when analyzing complex biological signals that are non-stationary and time varying. Wavelet transform methods, on the other hand, permit resolution of the time of event occurrences and changes in the frequency relationship over time. Signal processing by wavelet transform generates coefficients that represent the best-fit as a selected “mother wavelet” function is scaled and shifted along the source signal(21). There are two kinds of wavelet transforms, Discrete and Continuous wavelet transforms. With Discrete Wavelet Transform (DWT), the signal is decomposed into discrete frequency bands, without overlap of the time-frequency windows of the wavelet function. To detect major transitions that may be hidden in the noise of a physiological signal, the Maximal Overlap Discrete Wavelet Transform (MODWT)(21) can be employed. MODWT decomposes the signal into finer and finer frequency levels. As the level increases, large-scale approximations of the signal are obtained, and lower frequency components of the signal are well-resolved. MODWT of a signal allows for multi-resolution analysis (MRA) that reconstructs the decomposed time series as a sum of several new series that are aligned in time with the original signal. MODWTNMRA effects a zero-phase filtering of the signal. Features are time-aligned, unlike MODWT alone. Continuous Wavelet Transform (CWT) involves transformation of the signal by continuously changing the scaling and shifting factors. Although this introduces some information redundancy, it presents a more detailed, high resolution view of the characteristics of the signal. Coefficients generated by CWT are represented by a scalogram that is a visual representation of the frequency components of the signal as they change over time. In our experiments on cardiomyocytes loaded with the potentiometric fluorophore tetramethylrhodamine methyl ester (TMRM) and subjected to an in vitro I/R protocol, we used MODWTNMRA to identify the timing of the major ΔΨ_m_ depolarization during ischemia for each cell. CWT was utilized to analyze the more complex time varying frequency components of the ΔΨ_m_ oscillations observed in individual clusters of mitochondria during reperfusion. The image-processing and wavelet transform workflow, along with feature extraction from the images and scalograms obtained, allowed us to precisely determine the following: 1) the time point of ΔΨm loss for each cell during Ischemia, 2) the incidence of ΔΨ_m_ oscillation for each mitochondrial cluster and its frequency throughout the reperfusion period, 3) whether ΔΨ_m_ stabilized or irreversibly collapsed during reperfusion, and 4) the size distribution of the oscillating mitochondrial clusters.

#### (i) Identification of transition time-points of inner mitochondrial membrane potentials during Ischemia

To analyze a time-series of the mitochondrial inner membrane potential, we used MODWTNMRA to identify time-localized changes in the TMRM signal (using MATLAB’s signal processing toolbox). The TMRM signal from each cell during the Ischemic period was transformed with a sym4 wavelet with four levels of decomposition. Lower level decompositions involve higher frequencies and higher-level decompositions involve slower frequencies. For example, Fig.1 shows a raw TMRM signal from a single cell (A) decomposed into 4 levels using a sym4 wavelet transform (Fig. 1B). Level 1 has the frequency components between 0.033-0.017Hz, level 2 has 0.017-0.008 Hz, level 3 has 0.008-0.004 and level 4 has 0.004-0.002. All levels of decompositions have associated relative energies. For our purpose of estimating the ΔΨ_m_ depolarization time, we removed all higher frequency components with lower relative energy and reconstructed the signal by retaining the highest relative energy (of more than 99%) (Fig. 1C). We essentially filter out the ‘noise’ by this process. With this time-aligned reconstructed signal, we used the MATLAB function ‘findchangepoints’ to obtain the time point at which the reconstructed signal changed significantly (Fig. 1C). Time point of Ischemia depolarization can thus be automatically determined for several cells (Fig 1 E).

**Figure 1:**
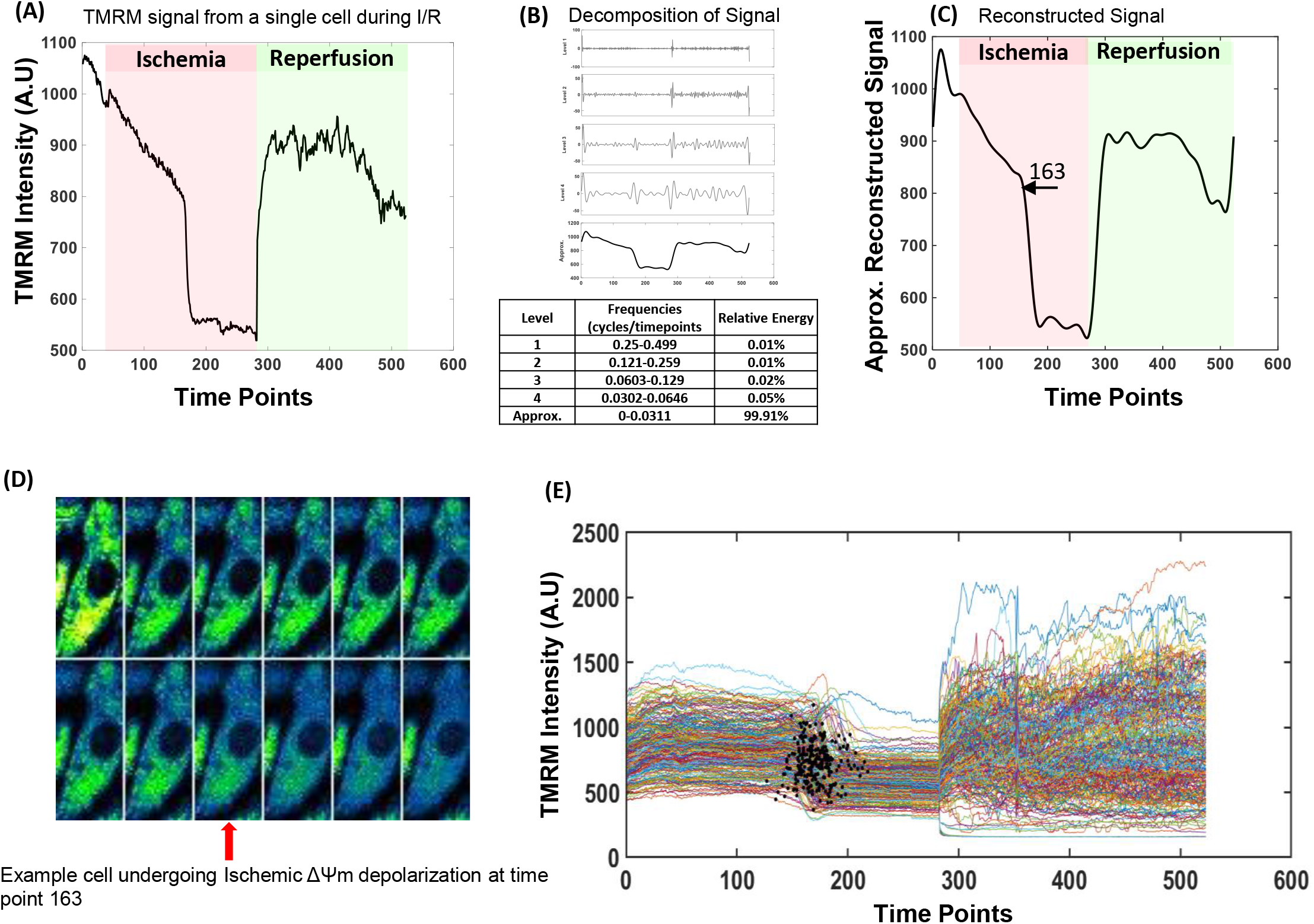
Estimating ΔΨ_m_ depolarization time during Ischemia by Multiresolution Signal Analysis. Identification of Ischemic depolarization time. Raw TMRM signals during Ischemic period (A) are decomposed using Maximal Overlap Discrete Wavelet Transform (B) and reconstructed by retaining the signal with the highest relative energy (C). MATLAB’s ‘findchangepoints’ function identifies the time point at which the signal changed significantly during Ischemia. Here, it is at time point 163. D) Example of a cell with the first image at baseline and subsequent images in the last phases of depolarization. Images are 15 seconds apart. TMRM intensity is abruptly diminished at time point 163. E) Example of Ischemia/ Reperfusion experiment with >100 cells where the black dots represent Ischemic depolarization time points.

#### (ii) Obtaining features and frequency components of ΔΨm oscillations during reperfusion

Mitochondria exhibited non-stationary oscillatory behavior throughout reperfusion (Fig 2). We categorized ΔΨ_m_ oscillatory behavior based on our visual observations of 10 experiments. There were five outcomes that were observed based on the oscillatory state of ΔΨ_m_ throughout the reperfusion time period, i.e., (i) ΔΨ_m_ oscillations persisting throughout, (ii) No or very few ΔΨ_m_ oscillations, (iii) ΔΨ_m_ oscillations that stabilized after oscillating initially, (iv) ΔΨ_m_ oscillations that occurred, but there was early ΔΨ_m_ loss, and (v) No ΔΨ_m_ oscillations occurred, and there was early ΔΨm loss (Fig. 2). We used a continuous wavelet transform (sym8) (in MATLAB’s signal processing toolbox), to process the TMRM signal and observed that the signal processing tool readily detected transitions and frequencies depicting the behavior of mitochondrial ΔΨ_m_ changes. Figure 2, right panel, shows the scalograms obtained after performing a wavelet transform of the TMRM signal. We observed that an oscillating cluster has high coefficients concentrated in the scale of ~ 3 to 10, corresponding to a frequency of 4.3-45mHz, which does not exist in the scalogram of the non-oscillating cluster or during Ischemia.

**Figure 2:**
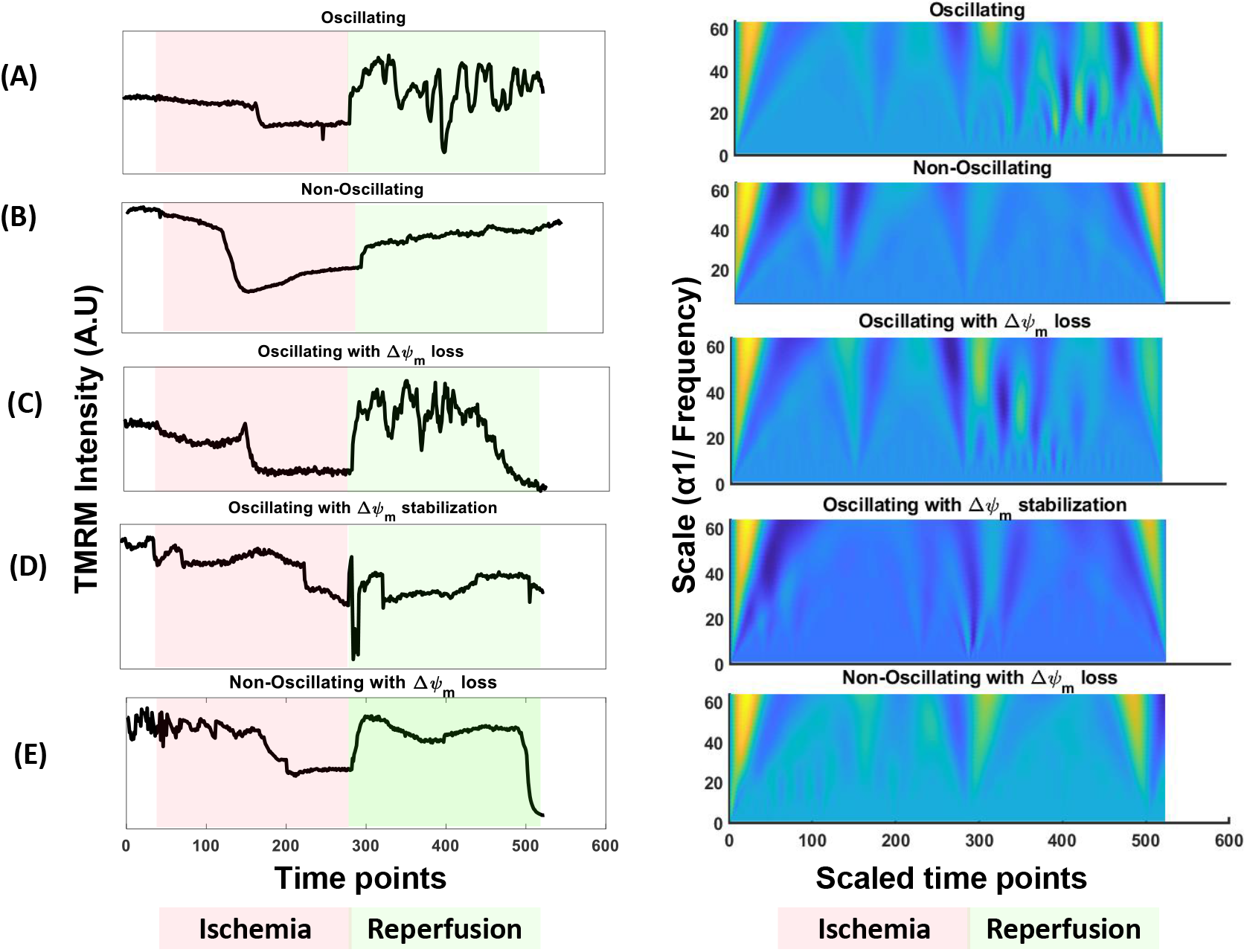
Mitochondria exhibit different kinds of ΔΨ_m_ oscillatory behavior during reperfusion. Mitochondria exhibit different ΔΨm oscillatory behaviors upon reperfusion. ΔΨm signals (TMRM fluorescence) for representative mitochondrial clusters during ischemia and reperfusion are shown in the lefthand panels and corresponding continuous wavelet transforms are shown in the righthand panels as scalograms. Frequencies and corresponding transition time points can be extracted from the scalograms. Mean TMRM Fluorescence intensities of an oscillating cluster (A), a non-oscillating cluster (B), an oscillating cluster exhibiting ΔΨm loss (C), a cluster that oscillates before stabilizing (D,) and a non-oscillating cluster exhibiting ΔΨm loss (E). The oscillating cluster (A) has high coefficients concentrated in the scale of ~ 1 to 10, corresponding to a frequency of 4.3-45mHz, which is absent in the scalogram of a non-oscillating cluster or during Ischemia.

This wavelet tool was then applied to detect transitions, frequencies and times associated with these changes automatically for a large number of cells (>100 per experiment) and mitochondrial clusters (> 400 per experiment). MATLAB/ FIJI platform was used to perform feature extraction for ΔΨ_m_ changes throughout the reperfusion time period (Figure 3). The procedure involved the following steps: (A) *image acquisition* with a confocal microscope using TMRM to monitor ΔΨ_m_ changes; (B) *cellular segmentation* using custom-made FIJI macros to separate each cell. The same thresholding method was applied to every image to outline each cell in the field of view; (C) By applying the threshold, each cell was separated into an image series; (D) *creation of an image Differential Stack* of the reperfusion phase of the image series by subtracting the n^th^ image from the (n-1)^th^ image. The sum of differentials in this stack could then be used to highlight the mitochondrial clusters that oscillate during the reperfusion period; (E) thresholding the z-projection of this differential image stack to obtain Regions of Interest (ROI) *outlining oscillating mitochondrial clusters*; (F) application of the ROIs to the reperfusion phase to *obtain TMRM signals for each cluster* through this time period; (G) *continuous wavelet transform of the TMRM signal* (with a sym 8 wavelet) to generate a coefficient matrix, visualized as a scalogram. The regions on the scalogram with large coefficients indicate where the mother wavelet fits the signal well. The x-axis represents the scaled time points and y-axis represents the scale (scale α 1/ frequency). Usually an oscillating mitochondrion shows high coefficient peaks corresponding to the scale range from 3-10. ΔΨ_m_ can also undergo larger transitions throughout reperfusion and these changes are reflected in the scalograms as high coefficient peaks; (H) importation of the resulting coefficient matrix as a scalogram-image and *extraction of predominant frequency features* as a function of reperfusion time. X and Y co-ordinates of the outlined maximum coefficients were obtained. The X-axis of the scalogram represents the time and the Y-axis, the scale (scale α 1/ frequency); (I) Mitochondrial oscillators associated with time are *classified into high/low frequency bands*. If a mitochondrial cluster oscillates in a particular frequency band at multiple times during reperfusion phase, then, an average of the frequency and the time is obtained. Thus, patterns of oscillatory behavior are obtained. We will henceforth refer to this routine as the MitoWave Analysis.

**Figure 3:**
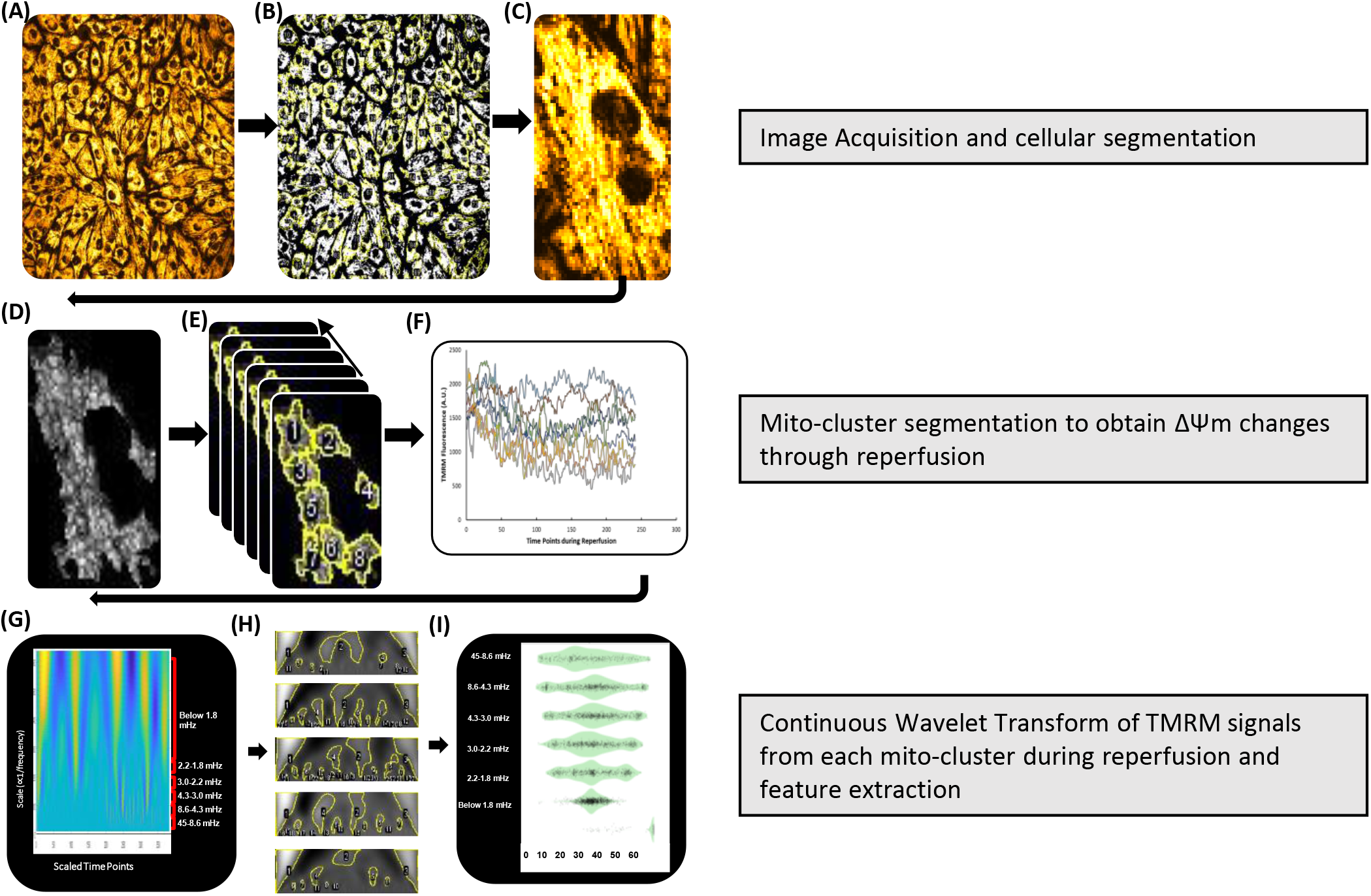
Workflow for characterizing ΔΨm oscillatory behavior during reperfusion: MitoWave. Schematic of Mito-wave analysis for ΔΨm feature extraction. It involves the following steps: A) Image Acquisition with a Confocal microscope using TMRM to monitor ΔΨm changes, B-C) Cellular Segmentation using custom-made FIJI macros to separate each cell, D) Differential stack Z-projection image for each cell is used to identify mitochondrial clusters that oscillate (MATLAB/FIJI Routine), E-F) TMRM fluorescence time course for each cluster is obtained, G) Scalograms are generated by continuous wavelet transform of the TMRM signal, H) Features and frequency components are extracted from the scalograms, and I) Mitochondrial oscillators are classified into high/low frequency bands to obtain patterns of oscillatory behavior as a function of reperfusion time.

(See supplement for ImageJ macros and MATLAB codes or on GitHub https://github.com/dashok1/MitoWave/releases/tag/v1.0.2).

## Results

### Defining oscillatory behavior patterns during Reperfusion

The behavior of each mitochondrial cluster was plotted into its corresponding frequency band, which varied over the reperfusion time period, represented as violin plots (Fig. 4). Frequencies were categorized as high frequency, ranging from 45 to 4.3 mHz (~22 seconds to 230 seconds), moderately fast frequencies ranging from 4.3-2.2 mHz (~ 230 seconds to 450 seconds), slow frequencies ranging from 2.2mHz to 1.8 mHz (~ 450 seconds to ~ 550 seconds) and below 1.8 mHz. Mitochondrial oscillators typically were present in the 45-4.3 mHz band. We also plotted the time at which there was complete ΔΨm loss during the reperfusion period. Applying Mito-Wave Analysis on ten in-vitro Ischemia/ Reperfusion experiments, we verified that our visual observations matched the quantitative analysis. In experiments where the mitochondrial oscillations persisted throughout the reperfusion period, high-frequency oscillators appeared at all time periods in the violin plots (Fig 4A) and when mitochondria had few/ no oscillations, the presence of high-frequency oscillators tapered off near 20 minutes of reperfusion (Fig 4B). We also observed, in some experiments, that mitochondrial oscillations occurred in the beginning of reperfusion, but started losing their ΔΨm during mid-late reperfusion, so the high-frequency oscillations tapered off, but shows up in the band where there is ΔΨm loss (fig 4C). We also observed in some experiments (Fig 4D), mitochondria exhibited few low amplitude or no oscillations at the beginning of reperfusion, so the number of high-frequency oscillators taper off around 20 minutes (similar to the distribution pattern of high frequency oscillators in fig 4B), but they begin to lose their ΔΨm around 25 minutes of reperfusion. Finally, in some experiments we observed that mitochondria stabilize their ΔΨm oscillations throughout the reperfusion time period (Fig 4E) where the presence of high-frequency oscillators taper off while ΔΨ_m_ is maintained during reperfusion. We then classified these experiments into the 5 different oscillation categories: Oscillating(4A), Non-Oscillating(4B), Oscillating with early ΔΨ_m_ loss(4C), Oscillating with early ΔΨ_m_ stabilization (4D) and Non-Oscillating with early ΔΨ_m_ loss (4E).

**Figure 4:**
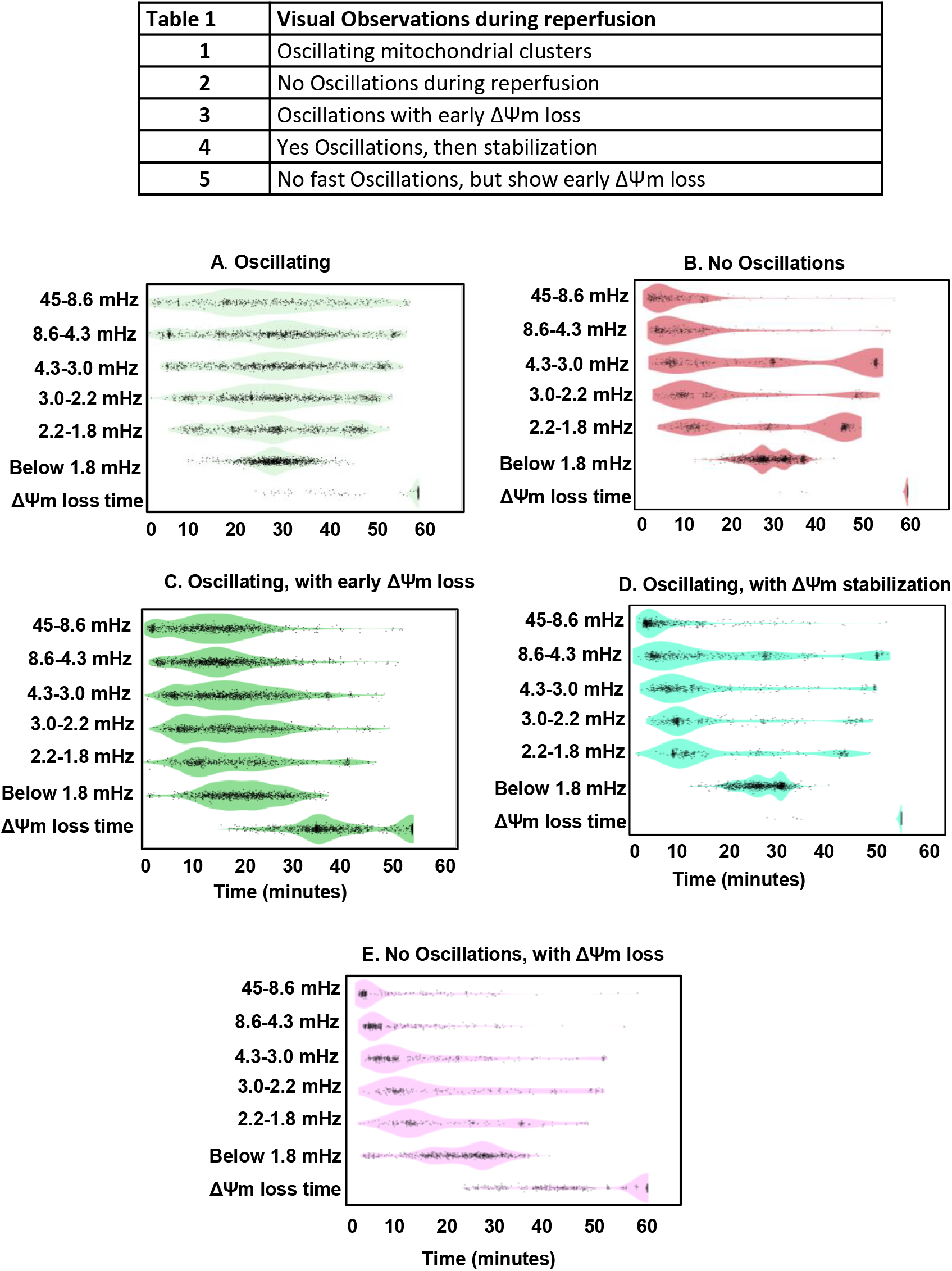
Defining ΔΨm Oscillatory patterns during reperfusion qualitatively and quantitatively: Visual observations of the Ischemia/Reperfusion image stack can qualitatively classify oscillatory behavior patterns of mitochondrial clusters during reperfusion. We classified oscillatory patterns from 10 experiments into 5 groups: Oscillating, Not Oscillating, Oscillating with early ΔΨm loss, Oscillating with ΔΨm stabilization, and Non-Oscillating clusters with Early ΔΨm loss (Table 1). By subjecting the TMRM signal from each mitochondrial cluster to MitoWave Analysis, we characterize oscillatory behavior quantitatively with violin plots (Fig. 4A-E). Each dot represents a mitochondrial cluster oscillating at a certain frequency corresponding to a certain time point. Visual observations (Table 1) are corroborated by results from the quantitative MitoWave analysis routine (fig 4A-E). We see that a mitochondrial cluster can change its oscillatory pattern throughout the reperfusion period, i.e., its frequency may change from one frequency band to another. Y-axis shows six frequency bands, as well as the time at which a mitochondrial cluster completely loses ΔΨm during reperfusion. X-axis represents the time of reperfusion.

### Predominant frequencies of mitochondrial clusters

We obtained the predominant frequencies of mitochondrial clusters by considering the first, fast frequency band (8.6-45mHz). If the mitochondrial cluster did not have a frequency in that band, the next frequency band was considered, and so on till the slowest frequency band. This way we could extract the frequencies that most closely represented mitochondrial oscillating frequencies. An average or a weighted average could be used since most mitochondrial clusters also have slow frequency components, but not all mitochondria have fast frequency components. Oscillating clusters have a frequency of 8.73±4.35mHz (1081 clusters), Non-Oscillating Clusters have 3.13±2.61mHz (732 clusters), Oscillating clusters with early ΔΨ loss have 9.56±3.66mHz (1402 clusters), Oscillating clusters with ΔΨ_m_ stabilization have 8.81±6.03mHz (1009 clusters) and Non-Oscillating clusters with Early ΔΨm loss have 6.82±4.63mHz (880 clusters) (figure 5A).

**Figure 5:**
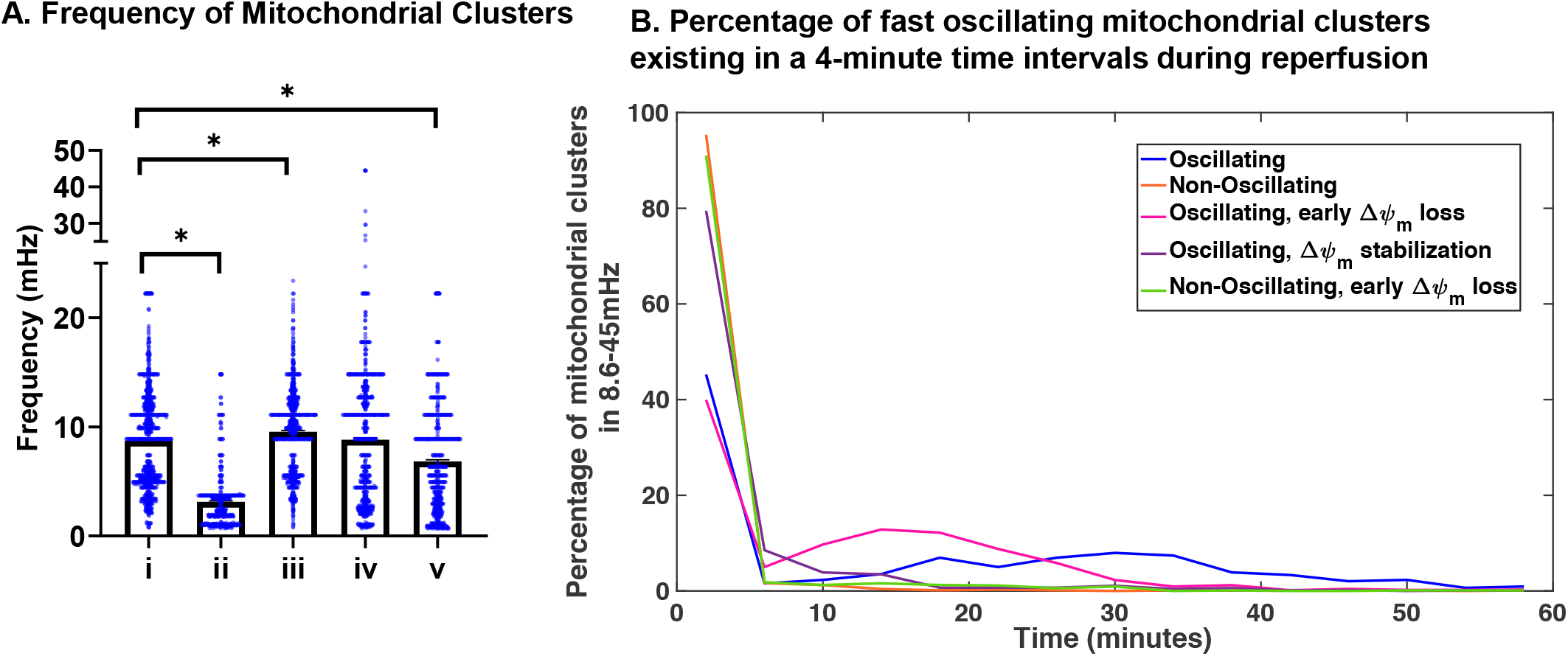
Predominant frequencies exhibited by mitochondrial clusters during reperfusion. The predominant frequencies exhibited by mitochondrial clusters fell within the 8.6 to 45mHz band. **A)** The mean predominant frequency ± SEM for i) Oscillating clusters, 8.73±4.35mHz(1081 clusters); ii) Non-Oscillating Clusters, 3.13±2.61mHz (732 clusters); iii) Oscillating cluster with early ΔΨ loss, 9.56±3.66mHz (1402 clusters); iv) Oscillating cluster with ΔΨ stabilization, 8.81±6.03mHz (1009 clusters); and v) Non-Oscillating clusters with early ΔΨm loss, 6.82±4.63mHz (880 clusters). One-way ANOVA was performed to determine statistical significance, * p <0.0001. **B)** Percentage of mitochondrial clusters oscillating in the 8.6-45mHz frequency band binned at 4-minute intervals during the reperfusion period.

Further, we analyzed the distribution of high frequency oscillators (in the 8.6-45mHz frequency band) to see how they vary throughout reperfusion time among the different categories. Clusters that didn’t have a frequency in this band (of 8.6-45mHz) were given a value of 0. We plotted the percentage of the different categories of oscillating clusters against time (5B). We observed that among the Oscillating category (blue line), 8-12% of mitochondria exhibited this high-frequency oscillations from 15-40 minutes of reperfusion. This was absent in the Non-Oscillating (orange line), in the Oscillating with early ΔΨ_m_ stabilization (Violet line) and the Non-Oscillating with ΔΨ_m_ loss (green line) categories. The Oscillating with early ΔΨ_m_ loss (pink line) shows ~ 7-15% of mitochondria exhibit high frequency only in the early reperfusion phase, till about 25 minutes, after which they do not. Further, we also statistically analyzed the distribution of these high frequency oscillators. A Kolmogorov-Smirnov non-parametric two sample test (kstest2 on MATLAB) was performed to test the null hypothesis that distribution of various oscillation behaviors were not different during the reperfusion time period. KS-test show significant differences between the different categories, comparing Oscillating and Non-scillating clusters, Oscillating and Oscillating with early ΔΨm loss, Oscillating and ΔΨm stabilizing clusters, and Oscillating and Non-scillating with early ΔΨm loss (p<0.0001). Thus, we quantitatively confirm our visual observations that the distribution of oscillating mitochondrial clusters that change dynamically over time are different between different categories of oscillating experiments.

### Frequency and mitochondrial cluster size are negatively correlated

We observed that in experiments where there were no/ few oscillations, mitochondrial clusters seem larger than in experiments where mitochondria had persistent oscillations. Previous reports in adult cardiac myocytes also showed that larger clusters have slower oscillations (11). Therefore, we wanted to check if this was true in Neonatal Cardiac myocytes as well. The Mito-Wave analysis of NMVMs subjected to I/R agreed with our visual observations. Oscillating mitochondria had the lowest area of 49.3μm^2^ vs a larger area of 65.92μm^2^ for non-oscillating mitochondria (Fig 6A). We performed non-parametric correlation coefficient analysis to understand the relationship between the size of mitochondrial clusters and its frequency. We found that there is a negative correlation between oscillating frequency and the size of the mitochondrial cluster, with a correlation coefficient of r= −0.58 (Fig. 6B). Mitochondrial cluster size decreased by ~4.56μm^2^ for every millihertz increase. This suggests that if mitochondria are organized in larger clusters, they undergo slower oscillations and may eventually stabilize ΔΨ_m_ and be protected against ΔΨm loss during reperfusion after Ischemia.

**Figure 6:**
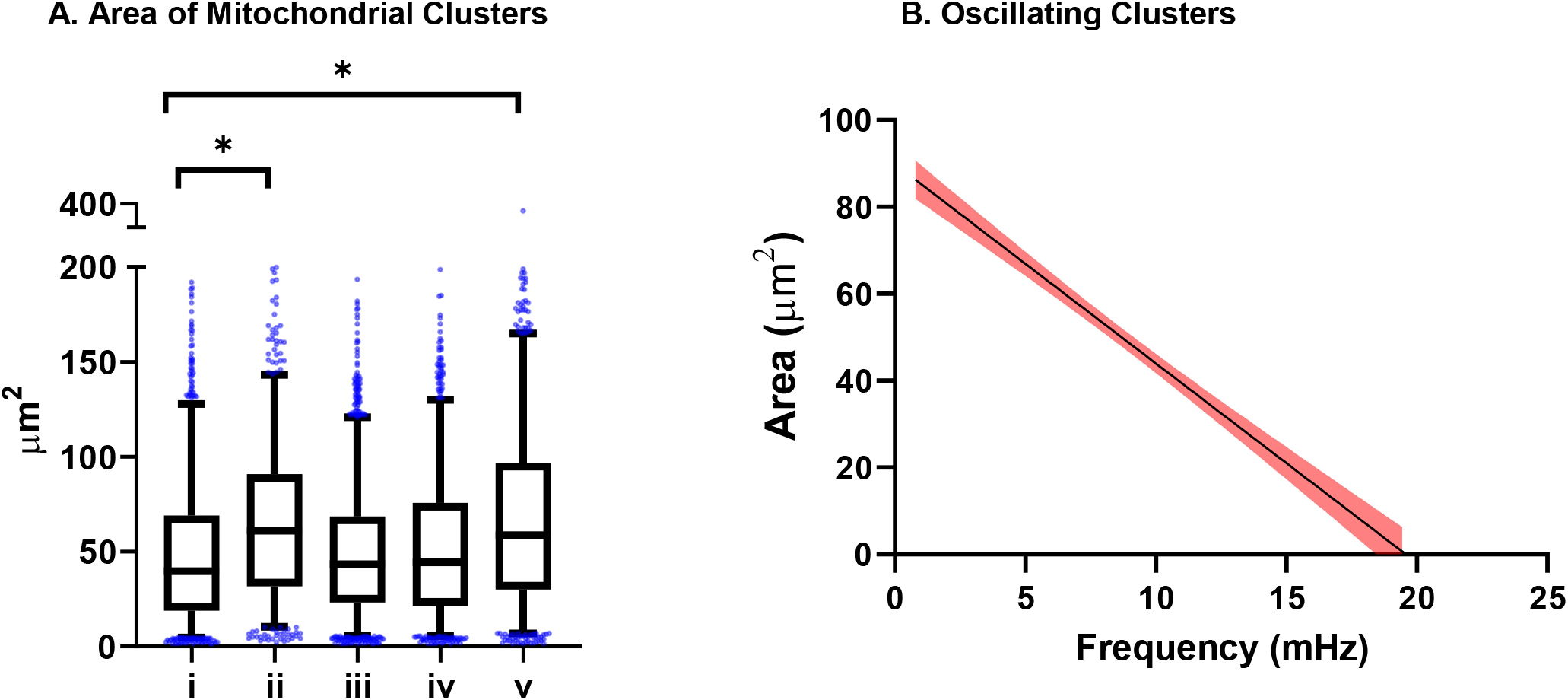
Mitochondrial Cluster size and Frequency relationship. **A)** Mitochondrial Cluster size and Frequency relationship. Areas of mitochondrial clusters were compared for clusters exhibiting different oscillatory behaviors (across several experiments). (i) Oscillating clusters had an area of 49.78±40.64μm^2^ (1081 clusters); (ii) Non-Oscillating Clusters, 65.97±42.07 μm^2^ (732 clusters); (iii) Oscillating cluster with early ΔΨ loss, 49.65±34.35μm^2^ (1402 clusters); (iv) Oscillating cluster with ΔΨ stabilization, 53.15± 39.38μm^2^ (1009 clusters); and (v) Non-Oscillating clusters with Early ΔΨm loss, 67.92±49.12μm^2^ (880 clusters). One-way ANOVA was performed to determine statistical significance, * p <0.0001. **B)** Frequency and cluster size show an inverse relationship. In Oscillating clusters, the area of the cluster decreases by ~4.56μm^2^ for every millihertz increase. 95% confidence intervals are plotted (red) with linear regression line (black).

### Time taken for ΔΨm loss during Ischemia and Reperfusion

Time to ΔΨ_m_ loss (reversible during Ischemia and irreversible during reperfusion) is an important indicator for mitochondrial resistance to instability during reperfusion after ischemia. It helps to understand if interventions to prevent mitochondrial instability and hence reperfusion injury are effective. We quantified the time to ΔΨ_m_ loss during Ischemia per cell (7A). The Oscillating category had a mean of 43.52±5.87 minutes to ΔΨ_m_ loss (i), Non-Oscillating took 46.36±9.17 minutes(ii), Oscillating with early ΔΨm loss took 35.62±9.25 minutes(iii), Oscillating with ΔΨm stabilization took 52.84±11.17 minutes (iv) and Non-Oscillating clusters with Early ΔΨ_m_ loss took 30.46±7.81 minutes(v). We also quantified the time to ΔΨ_m_ loss per mitochondrion during reperfusion (7B). We plotted the time against the percentage of mitochondria. Oscillating clusters take 58.71± 4.75 minutes to lose ΔΨ_m_; Non-Oscillating clusters did not lose their ΔΨm till the end of reperfusion at 60.25 minutes; Oscillating clusters with early ΔΨm loss take 45.8±11.05 minutes; Oscillating clusters with ΔΨm stabilization take 59.66±3.96 minutes and Non-Oscillating clusters with early ΔΨm loss take 53.38± 10.99 minutes.

### Correlation between Ischemic depolarization time point and ΔΨm oscillation frequency

We wanted to understand if there was any link between mitochondrial recovery during reperfusion and the time to ΔΨ_m_ loss during Ischemia. We compared the empirical cumulative distribution functions between different oscillation categories during Ischemia (8A) and reperfusion (8B). We found that late ΔΨ_m_ loss during Ischemia correlated with mitochondrial ΔΨ_m_ stabilization during reperfusion.

## Discussion

Over the course of ischemia-reperfusion, the mitochondrial networks of cultured neonatal mouse cardiomyocytes displayed complex spatiotemporal patterns, including bistability and time-varying oscillatory behavior, presenting significant challenges to analysis. The present work combined image segmentation with the versatility of wavelet transforms to quantify key transitions associated with the pathophysiology of I/R injury in an unbiased manner. Essential information could be captured in a semi-automated workflow, including the time-to-mitochondrial depolarization during ischemia, frequency of ΔΨ_m_ oscillation of individual mitochondrial clusters upon reperfusion, and time to catastrophic loss of ΔΨ_m_ with prolonged reperfusion. Subsequent data reduction permits one to make statistical comparisons between different experiments to determine if a given treatment or intervention has significant effect on mitochondrial function (Fig 3).

We have previously reported that adult cardiomyocytes subjected to metabolic or oxidative stress undergo spontaneous oscillations in ΔΨ_m_ that occur either in small clusters or are synchronized across the whole cell (11). Cell wide ΔΨ_m_ synchronization is observed after a critical number of mitochondria in the network show oxidative stress, a phenomenon we termed “mitochondrial criticality”(22). Synchronization of mitochondria in the organized array of the adult myocyte depends on ROS-dependent neighbor-neighbor interactions between organelles, with long range cluster interactions following the behavior of a percolation lattice(23). In neonatal myocytes, the mitochondrial network is less ordered and reperfusion-induced oscillations are less likely to be synchronized throughout the entire network (13), consistent with a short effective diffusion distance for ROS-induced ROS release(10). In contrast, when the system is forced by a uniform environmental stress, such as ischemia, mitochondrial network depolarization occurs on a cell-by-cell basis, likely determined by the anaerobic ATP-generating capacity and glycogen store of the individual cells. The average time to ischemic ΔΨ_m_ depolarization for a given coverslip was compared to the oscillatory behavior of mitochondrial clusters on reperfusion (Fig. 7 & 8). Interestingly, early ΔΨ_m_ loss during ischemia correlated with early ΔΨ_m_ loss during reperfusion; however, this was equally true for both oscillating and non-oscillating clusters, suggesting that there is no specific protective advantage of the oscillatory behavior. In fact, there was a trend towards earlier depolarization during reperfusion for oscillating versus non-oscillating mitochondrial clusters. At least concerning mitochondrial recovery after reperfusion, these findings argue against the idea that oscillations in metabolism might preserve a higher average ATP/ADP ratio while decreasing free energy dissipation compared to steady state operation (24). Instead, mitochondrial ΔΨ_m_ oscillation could simply be an inevitable consequence of the nonlinear control properties of the nonlinear bioenergetic system. In addition, late ΔΨ_m_ loss during ischemia correlated with ΔΨ_m_ stabilization after oscillation on reperfusion. Together these data indicate that mitochondrial energetic recovery strongly depends on resistance to initial ischemic depolarization, consistent with data from intact perfused hearts(25).

**Figure 7:**
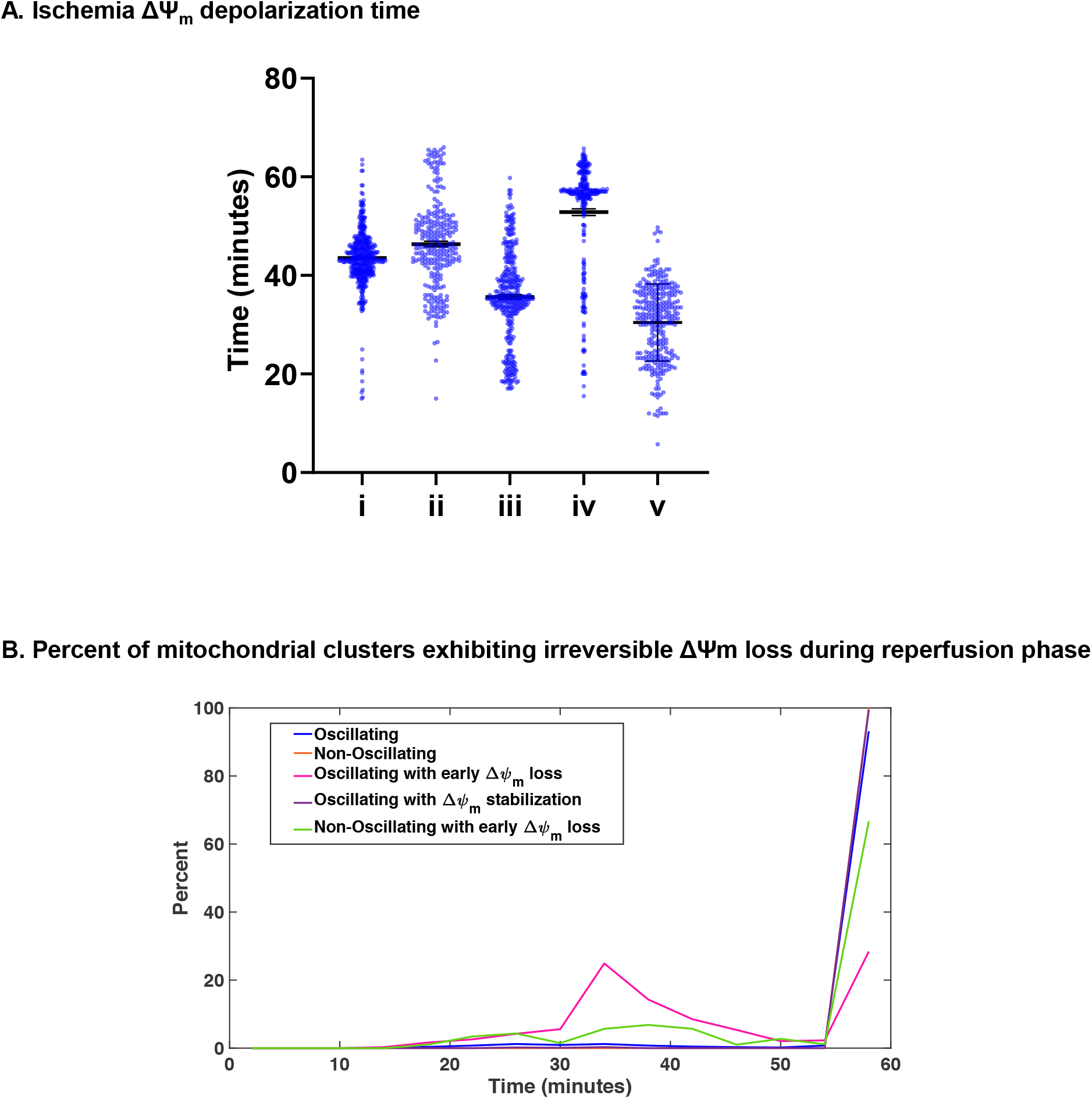
Time taken for ΔΨ_m_ loss during Ischemia and Reperfusion. **A)** Time taken for ΔΨ_m_ loss during Ischemia versus the ensuing oscillatory behavior on reperfusion. (i) Oscillating clusters maintained ΔΨ_m_ until 43.52 ±5.87 minutes; (ii) Non-Oscillating clusters, 46.36±9.17 minutes; (iii) Oscillating with early ΔΨ_m_ loss, 35.62±9.25 minutes (iv) Oscillating with early ΔΨ_m_ stabilization, 52.84±11.17 minutes and (v) Non-Oscillating clusters with Early ΔΨ_m_ loss, 30.46±7.81 minutes. **B)** Percentage of mitochondrial clusters exhibiting irreversible ΔΨ_m_ loss during reperfusion. Oscillating clusters lost ΔΨ_m_ on average at 58.71± 4.75 minutes of reperfusion; Non-Oscillating clusters maintained stable ΔΨ_m_ to the end of 60.25 minutes of reperfusion; Oscillating clusters with early ΔΨ_m_ loss depolarized at 45.8±11.05 minutes; Oscillating clusters with ΔΨ_m_ stabilization lasted 59.66±3.96 minutes, and Non-Oscillating clusters with early ΔΨ_m_ loss depolarized at 53.38± 10.99 minutes

**Figure 8:**
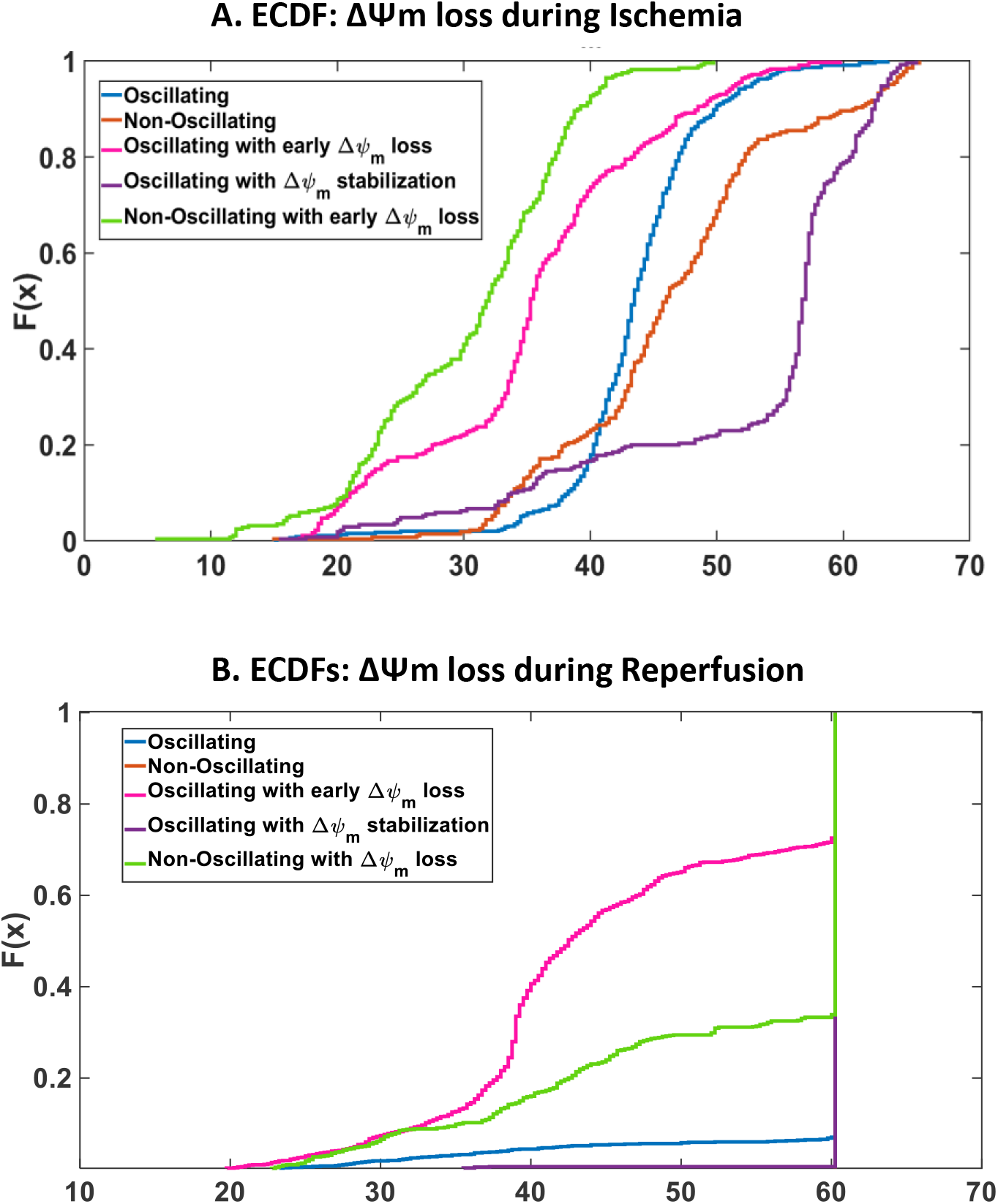
Relationship between Ischemia ΔΨ_m_ depolarization time with Oscillatory behavior during Reperfusion. Relationship between Ischemic ΔΨm depolarization time and Oscillatory behavior during Reperfusion **A)** Empirical Cumulative Distribution functions showing the probability of depolarization (F(x)) as a function of time (x) during Ischemia. **B)** Empirical Cumulative Distribution functions showing the probability of depolarization (F(x)) as a function of time (x) during Reperfusion. Mitochondrial stabilization during reperfusion correlated with late ΔΨ_m_ loss during Ischemia (purple line).

The present findings show that in NMVMs subjected to I/R, ΔΨ_m_ oscillation frequency is inversely correlated with cluster size (Fig. 6). This is in agreement with the negative correlation obtained by wavelet transform analysis of adult myocytes under oxidative stress, with large mitochondrial clusters showing slow ΔΨ_m_ oscillations that could span the entire cell with a stereotypical frequency of 1 - 10 mHz(11). Synchronization of a network of dynamically coupled oscillators spanning a broad frequency range to a single dominant frequency is common to physical, chemical and biological systems. The lack of synchronization in NMVMs and the broader frequency distribution (Fig. 5) may be the result of the more disorganized arrangement of mitochondria in neonatal myocytes or weaker coupling between mitochondria in the immature cells.

The method described here provides a way to uncover and quantify different mitochondrial responses to I/R stress that might otherwise be overlooked if one were to only examine the average behavior of a monolayer, of individual cells, or at single time points during a protocol (e.g., measuring lactate dehydrogenase release as an index of damage after reperfusion). A current limitation of the method is that it would be affected by significant movement of the objects being analyzed in the optical field, which was minimal in our experiments. In the future, it might be possible to further develop the approach by incorporating object tracking methods. Nevertheless, the approach is applicable to any spatially-distributed system of time varying oscillatory signals. Unlike Fourier transform analysis, the underlying oscillator frequencies and phases do not have to be time invariant and the method is largely immune to changes in signal offset (such as photobleaching) or background artifacts. This novel approach, which standardizes the quantitative analysis of complex biological signals, opens the door to in depth screening of the genes, proteins and mechanisms underlying metabolic recovery after ischemia-reperfusion.

## Acknowledgements

We are grateful to Dr. Amitabh Basu, for discussions on statistical analysis and MATLAB coding This work was supported by NIH grants R01HL137259, R01HL134821 (BO’R) and F31HL134198 (DA).

## Author Contributions

D.A. performed experiments, performed analysis, wrote codes for analysis and wrote the paper. B.O’R conceived the idea and wrote the paper

## Supplemental information

1. MitoWave Analysis Routine ImageJ and MATLAB codes (also on GitHub https://github.com/dashok1/MitoWave/releases/tag/v1.0.2 and zenodo https://doi.org/10.5281/zenodo.3820382)
2. *In-vitro* Ischemia/Reperfusion experiment movie showing mitochondria Oscillating throughout reperfusion: Oscillating Category
3. *In-vitro* Ischemia/Reperfusion experiment movie showing mitochondria Not Oscillating throughout reperfusion: Non-Oscillating Category
4. *In-vitro* Ischemia/Reperfusion experiment movie showing mitochondria Oscillating, then losing ΔΨ_m_ during reperfusion: Oscillating with early ΔΨ_m_ loss Category
5. *In-vitro* Ischemia/Reperfusion experiment movie showing mitochondria Oscillating, then stabilizing during reperfusion: Oscillating with early ΔΨ_m_ stabilization Category
6. *In-vitro* Ischemia/Reperfusion experiment movie showing mitochondria Not Oscillating and exhibiting early ΔΨ_m_ loss during reperfusion: Non-Oscillating with early ΔΨ_m_ loss Category

## Notes

**Conflict of interest:** The authors have no conflicts to disclose

